# From leaves to defenders: how the amount and dispersion of leaf damage affect extrafloral nectar production and ant-mediated protection in wild cotton

**DOI:** 10.1101/2025.07.31.667819

**Authors:** Víctor Hugo Ramírez-Delgado, Yeyson Briones-May, Alejandra Garnica-Cabrera, Juan Sánchez-Durán, Lucía Martín-Cacheda, Xoaquín Moreira, Ted C J Turlings, Miguel Vásquez-Bolaños, Luis Abdala-Roberts

## Abstract

1. Extrafloral nectar (EFN) drives mutualistic interactions between plants and ants. However, EFN production is costly, and its induction is predicted to vary with herbivore-related factors. We examine how the amount and within-plant damage spatial uniformity (i.e., dispersed and concentrated damage) affect EFN production and ant-mediated defence in wild cotton (*Gossypium hirsutum*). Previous work suggests that severe and concentrated damage is costliest to plants, leading us to hypothesise that such damage would induce greater EFN production and, consequently, stronger ant attraction and protection.
2. We conducted a greenhouse experiment in which plants were subjected to one of the following mechanical damage treatments: control (no damage), low damage (two damaged leaves, 30% of area removed each), high concentrated damage (two damaged leaves, 60% each), and high dispersed damage (four damaged leaves, 30% each; i.e., same total area than concentrated damage). The day after treatments, we measured EFN volume and sugar content from one nectary per plant (a per-nectary EFN response), and the number of secreting nectaries per plant (a proxy for whole-plant level EFN response). Subsequently, we performed a field experiment using the same treatments to evaluate EFN-mediated ant recruitment and ant-provided protection by placing caterpillars on plants.
3. Both high-damage treatments increased EFN volume per nectary, whereas low damage did not. However, against our prediction, dispersed damage was the only treatment that increased the number of secreting nectaries. Consistently, once placed in the field, plants with dispersed damage recruited more ants and caterpillars on them were more likely to be attacked by ants.
4. Contrary to expectations, plants subjected to dispersed damage exhibited a greater total investment in EFN, presumably via local induction of more nectaries per plant, and thereby enhancing ant-mediated defence. This study highlights the importance of herbivory intensity and within-plant spatial uniformity in shaping ant-plant interactions.

## Introduction

Extrafloral nectaries are specialised structures present in more than 100 families of plants (Weber & Keeler, 2012) that secrete a sugary, energy-rich solution. This secretion can occur constitutively (always present, i.e., basal levels) on leaves and stems but is also highly inducible (i.e., increased production over and above basal levels) in response to herbivory (Rico-Gray & Oliveira, 2008; Heil, 2015). Extrafloral nectar (EFN) serves as an important food source for ants and other insects, which in turn, often protect the plant by deterring or attacking herbivores (Heil & McKey, 2003; Rico-Gray & Oliveira, 2008; Rosumek et al., 2009). However, EFN induction is highly variable and context-dependent, shaped by both biotic and abiotic factors, and modulated by the balance of its benefits (via indirect defence) and costs to the plant across different ecological contexts (Bronstein, 1998; Bronstein et al., 2006; Kessler & Heil, 2011; Marazzi et al., 2013; Interian-Aguiñaga et al., 2025). Addressing this context dependency is key for understanding the ecological impacts and evolutionary stability of EFN-mediated mutualisms.

Multiple herbivory-related factors regulate EFN induced responses. Commonly, EFN induction increases in a dose-dependent manner as a function of the amount of herbivory (Rios et al., 2008; Xu et al., 2014; Huang et al., 2015; Calixto et al., 2021), suggesting that plants can fine-tune their defensive investment based on perceived herbivore pressure. Here studies have found that higher levels of leaf damage often trigger increased EFN production, which in turn attracts more ant defenders to help mitigate further herbivory (Heil et al., 2001; Ness, 2003; Nogueira, 2025). This relationship is not always linear, with some studies reporting threshold effects where EFN secretion significantly increases only after damage surpasses a certain level (Pimenta et al., 2025). Similar patterns have been shown for other plant defensive traits, such as chemical defences, where induction thresholds as well as saturation (no further increase in induction beyond a given damage level) can constrain plant induced responses (e.g. Underwood, 2010). These findings suggest that induction of plant defences, including EFN, operates within physiological or ecological limits. Within this context, research on EFN responses to varying levels of herbivory intensities has mounted in recent years (e.g., Xu et al., 2014; Abdala-Roberts et al., 2019b; Calixto et al., 2021; Interian-Aguiñaga et al., 2025; Pimenta et al., 2025), but further studies are needed to clarify how these patterns influence ant-plant interactions.

The spatial uniformity of herbivore damage within a plant can also play a crucial role in shaping plant fitness and the induction of defensive traits (Wetzel et al., 2023). Early work suggests that plants tolerate dispersed damage better than concentrated damage, implying higher fitness costs associated with the latter (Marquis, 1992; 1996; Mauricio et al., 1993). The reduced tolerance to concentrated damage has been proposed to stem from the modular nature of plants and physiological limitations for resource movement between modules. When damage is concentrated on specific branches (or leaves), resource limitations in those areas cannot be easily compensated for by other parts of the plant, leading to localised stress and reduced whole-plant level re-growth (Marquis, 1996). In accordance, Optimal Defence Theory predicts that plants should induce stronger defensive responses under concentrated damage compared to dispersed damage, in order to offset the greater cost associated with more localised herbivory (Holland et al., 2009; Moreira et al., 2012; Calixto et al., 2021). However, most research to date has focused on tolerance mechanisms associated with sectoriality (i.e. resource reallocation constraints between plant modules limiting regrowth or compensatory growth) whereas work examining how damage spatial uniformity affects the induction of resistance-related traits have been largely ignored. In one of the few exceptions. Ohnmeiss et al. (1997) found that concentrated damage increased nicotine levels (a direct defence) to a greater extent than dispersed damage in *Nicotiana sylvestris*. It has been hypothesised (Marquis, 1996) and shown empirically by at least one study that stronger induced resistance responses to concentrated damage (e.g., local induction) contribute to the spread of subsequent damage throughout the canopy (e.g., Karban & Yang, 2020), reducing subsequent herbivory costs. Alternatively, defensive responses could be modulated by damage thresholds within plant organs. Once damage exceeds critical thresholds, a plant may no longer invest in defending a compromised organ (Wetzel et al., 2023), and instead allocate more defences to less damaged or intact organs. These ideas remain largely untested, and, to our knowledge, no study to date has evaluated how within-plant damage spatial uniformity influences EFN induction and ant-plant interactions, or disentangled these effects from other herbivory-related factors (e.g., total damage amount, see above).

Here, we used wild cotton (*Gossypium hirsutum* L.; Malvaceae), an extrafloral nectary-bearing, myrmecophytic shrub distributed along the northern coastline of Yucatan (Mexico), to evaluate the effect of leaf damage amount and within-plant dispersion on the production of EFN. We also examined how these EFN induction patterns influence ant recruitment, as well as how effectively ants deter or prey upon herbivores. Previous work with wild cotton has shown that EFN induction enhances ant recruitment and defensive responses against herbivores (Vázquez-Barrios et al., 2021), suggesting that EFN can provide fitness benefits via indirect defence, though not universally (Reyes-Hernández et al., 2022). Wild cotton plants also show cotton plants are less able to compensate (in terms of regrowth) after high levels of damage as well as in response to concentrated (vs. dispersed) damage (Quijano-Medina et al., 2021, Ramírez-Delgado et al., 2025), implying higher fitness costs under these herbivory scenarios. Based on this, we hypothesised that high and concentrated damage would elicit stronger EFN induction (in accordance with expected higher costs due to these types of damage), leading to greater ant recruitment and attack rates on herbivores. Overall, by disentangling the effects of herbivory intensity and within-plant damage spatial uniformity on EFN induction and ant-plant interactions, this study provides novel insights into how herbivory-related factors may shape indirect plant defence.

## Methods

### Study species

Wild cotton (*Gossypium hirsutum* L.) is a coastal myrmecophytic shrub native to Mexico, Central America, and the Caribbean, that is particularly abundant in the Yucatan Peninsula (Coppens d’Eeckenbrugge & Lacape, 2014). Leaf chewers, such as grasshoppers and caterpillars, cause most of the damage to this species, with the specialist *Alabama argillacea* Hübner, 1823 (Erebidae) being particularly abundant and damaging at some sites (Nascimento et al., 2011). Leaf damage exhibits considerable variability—up to 3.5-fold difference among plants within populations, and up to three-fold variation in damage distribution across leaves within individual plants (Abdala-Roberts et al., 2019a). In addition, there is some evidence suggesting that high and concentrated damage are costlier for wild cotton plants (Quijano-Medina et al., 2019; Ramírez-Delgado et al., 2025). Wild cotton secretes EFN from specialised nectaries located along the mid-vein and sometimes on secondary veins. Several studies have shown that EFN production in wild cotton increases primarily in response to mechanical leaf damage (Abdala-Roberts et al., 2019b; Briones-May et al., 2023), with similar findings reported for cultivated cotton (Wäckers & Bonifay, 2004; Llandres et al., 2019). In turn, a diverse ant community is associated with wild cotton in situ, including species of *Camponotus*, *Pseudomyrmex, Monomorium, Pheidole Brachymyrmex*, *Dorymyrmex*, and *Crematogaster* (Vázquez-Barrios et al., 2021, Reyes-Hernandez et al., 2022). Many of these ant species consume EFN and can play a defensive role by deterring or consuming insect herbivores (Vázquez-Barrios et al., 2021). Also, in a previous work, Abdala-Roberts et al., (2019b) found that higher levels of damage more strongly induced EFN, but to date, the effects of within-plant damage spatial uniformity on EFN induction have not been tested or disentangled from the effects of damage amount, and the consequences of these damage patterns for ant attraction and ant-plant-herbivore interactions have not been investigated.

### Plant material and propagation

We collected seeds from seven adult plants (hereafter referred to as genotypes) located at a coastal scrubland site near the port of Sisal, Yucatán, Mexico (21°11’ 45.3” N, 89°57’ 18” W). We germinated seeds and transplanted them into 2-litre polyethene bags filled with a soil mixture of sand, native deciduous forest soil, and perlite (1:2:1). We maintained plants in a greenhouse at the Campus de Ciencias Biológicas y Agropecuarias of the Universidad Autónoma de Yucatán (20° 51’ 57” N, 89° 37’ 20” W), where they were watered three times per week with 300 ml of water. At the time the experiments were conducted, plants were four months old.

### Greenhouse experiment: treatment effects on extrafloral nectar

We used a total of 79 plants to evaluate the effects of leaf damage treatments on EFN induction. We placed plants 90 cm apart and subjected them to one of the following mechanical damage treatments using serrated scissors, with genotypes evenly represented across groups: a) control: no leaves were damaged—18 plants; b) low damage: one side was removed from each of two leaves (leaves 2 and 4, counting downward from the apical meristem)—20 plants; c) high concentrated damage: two sides were removed from each of two leaves (2 and 4)—22 plants; and d) high dispersed damage: one side was removed from each of four leaves (2 to 5)—19 plants. Damage treatments were applied over two consecutive days to mimic the gradual progression of damage from feeding patterns by insects, such as lepidopterans and orthopterans, which are the main herbivores for *G. hirsutum* (Abdala-Roberts et al., 2019a). We did not damage leaf 1 since it is often still under expansion and this may influence local induced responses compared to mature leaves. To standardize damage application, we first calibrated the technique using leaves from non-experimental plants of a similar age and verifying the percentage of leaf area removed by analysing photographs of these leaves analysed in ImageJ software (Schneider et al., 2012). On the first day, we removed 15% of the area per leaf by cutting one side of each of two or four leaves per plant for low and high dispersed damage treatments, respectively, or 15% of the area for both sides of each of two leaves per plant for the high concentrated damage treatment (30% of the total area of each leave). The following day, we removed an additional 15% from each previously damaged side of the same leaves. As a result, plants in the low and high dispersed damage treatments had two or four leaves damaged, respectively, with 30% area removed per leaf in both cases. In contrast, plants in the high concentrated damage treatment had two leaves damaged, each with 60% of the area removed. Thus, highly damaged plants experienced twice the total amount of damage compared to low-damaged plants. At the same time, the two high-damage treatments differed only in the spatial uniformity of damage, not in total damage amount (Fig. S1, supplementary material). Comparisons between the low- and high-damage treatments allowed us to test for an effect of damage amount, while the comparison between the two high-damage treatments (concentrated vs. dispersed) allowed us to test for an effect of within-plant damage spatial uniformity (Martín-Cacheda et al., 2025). It is important to note that in cotton, mechanical damage alone is sufficient to elicit EFN induction of a magnitude comparable to that caused by natural herbivory (Wäckers & Wunderlin, 1999). This includes induction levels similar to those observed under varying degrees of damage amount and within-plant damage dispersion caused by natural herbivory (Ramírez-Delgado et al., 2025). Thus, mechanical damage provides a realistic proxy for testing EFN defence induction, while offering greater precision and control over the amount and distribution of leaf damage (Abdala-Roberts et al., 2019b). In other instances, however, as for chemical defences, elicitors from insect oral secretions have been shown to influence cotton chemical defences (Arce et al., 2021).

On the morning following the completion of damage treatments, between 06:00 and 07:00, we collected EFN from leaf 2 of each plant using 5-μL capillary tubes (micropipettes Blaubrand ® intraMARK, colour code white, Germany). Twenty-four hours prior to nectar sampling, we removed nectar from all nectaries with wetted cotton, ensuring a baseline for EFN measurements during the sampling day. We selected leaf 2 as it was the most apical leaf with damage, and EFN induction is typically strongest in younger leaves (1-3) and at the site of damage (i.e., local induction; Abdala-Roberts et al., 2019b). After measuring EFN volume in the capillary tubes, we determined sugar concentration (° Brix) using a refractometer (Atago Master, Japan). We then calculated total sugar produced per sample as: total sugar = nectar volume × nectar sugar concentration (° Brix). One-degree Brix (1 ° Brix) is equivalent to 1 gram of sugar dissolved in 100 grams of solution (Kimball, 1991). Finally, we recorded the number of active (secreting) nectaries per leaf by visual inspection. To ensure that nectaries classified as inactive were truly not secreting, we gently probed each one with a capillary tube to confirm the absence of EFN. In addition to per-nectary responses reflecting local induction (volume, sugars), the number of secreting nectaries provided a measure of whole-plant response, involving not only local but likely also systemic induction (EFN produced by non-damaged leaves).

### Field experiment: treatment effects on ant recruitment and their anti-herbivore responses

A week after the greenhouse experiment, we transported a second group of 101 plants (same age as those used in the greenhouse EFN induction experiment) to coastal scrubland site near Sisal, Yucatan. We placed plants at nearby wild cotton plants from which seeds for the experiments were sourced. Plants were spaced one meter apart in an open area surrounded by natural vegetation. We assigned plants randomly to the same leaf damage treatments described above (control: 28 plants; low damage: 24 plants; high concentrated damage: 27 plants; high dispersed damage: 22 plants), using the exact same methodology, with genotypes evenly represented across treatment groups. Plant positions were also randomised. Throughout this field experiment, we watered the plants twice a week. Two days after placement, we applied the mechanical damage treatments. The following morning, we recorded ant abundance for 2 min per plant between 9:00 and 11:00 AM over two consecutive days. We collected ant specimens for taxonomic identification in the laboratory. The day after the second ant survey, we conducted ant attack bioassays by placing caterpillars over two consecutive days. For this, each morning at 9:00 AM, we placed a frozen third-or fourth-instar caterpillar of *A. argillacea* on each plant, gently securing it to an apical leaf with a metal clip. The use of dead caterpillars allowed us to standardise caterpillar position and eliminate behavioural and chemical variability associated with live specimens (Lövei & Ferrante, 2017; Nagy et al., 2020). We observed no saprophagous insects on the plants during the bioassays. Per day, and based on prior bioassays of this type, we inspected each plant 100 minutes after caterpillar placement to determine whether the caterpillar had been attacked (scored as 1) or not attacked (scored as 0). An attack was defined as any instance in which one or more ants made direct contact with the caterpillar and exhibited aggressive behaviour or attempted to remove it. Cases in which the caterpillar was missing for unknown reasons (< 6% of observations) were excluded from statistical analyses.

### Statistical analyses

We used generalised linear mixed models (GLMMs) to analyse nectar volume, total sugar production, the proportion of active nectaries (number of leaves with active nectaries relative to the total number of leaves per plant), the number of patrolling ants, and attack by ants. For both ant-related response variables, we pooled the counts across all observed ant species for increased power and to test for community-level ant responses. In all models, we included treatment as the main fixed effect and plant height as a covariate. We included plant genotype as a random effect to account for genetic or maternal effects. For the models analysing ant abundance and attack by ants, we also included survey day and plant ID as random factors to account for temporal variation and repeated measures taken on the same plants across sampling days, respectively. For nectar volume and total sugar production, we used a Tweedie distribution with a log link function, which is suitable for continuous, non-negative, and right-skewed data that include zeros (Hasan & Dunn, 2015). The proportion of active nectaries was analysed using a beta distribution with a logit link function, ant abundance (pooled across species was modelled using a negative binomial distribution with a log link function to account for overdispersion, while the probability of attack by ants was modelled using a binomial distribution with a logit link function which estimates the natural logarithm of the odds (log-odds) of an attack occurring. These log-odds were then back-transformed to provide an estimate of the probability of ant attack.

All statistical analyses were performed in R 4.4.2 (R Core Team, 2024). We used the “glmmTMB” package (Brooks et al., 2017) to fit the GLMMs, and the “DHARMa” package (Hartig, 2024) to evaluate models fit. To test for main effects, we employed the *Anova*() function from the “car” package (Fox & Weisberg, 2019), using a Wald χ^2^ tests. When treatment effects were significant, we used the “emmeans” package (Lenth, 2024) to perform Tukey pairwise comparisons between groups and to extract back-transformed estimated marginal means (EMMs) with their 95% confidence intervals (95% CIs). We visualised these results using the “ggplot2” package (Wickham, 2016). Model estimates are reported throughout the Results section as EMMs with 95% CI, in the format: EMM [2.5% lower CI, 97.5% upper CI].

## Results

### Extrafloral nectar

Results from the greenhouse experiment showed that leaf damage significantly influenced EFN volume (Table 1). Both high damage treatments—concentrated (0.589 [0.409, 0.850] µL) and dispersed (0.694 [0.479, 1.01] µL)—resulted in significantly more EFN volume produced compared to the control group (0.197 [0.107, 0.36] µL) (Fig. 1a). The low damage treatment (0.457 [0.3, 0.697] µL) did not differ significantly from either the control or the high damage treatments (Fig. 1a). Leaf damage also had a significant effect on total sugar production (Table 1). In this case, all damage treatments—high concentrated (0.140 [0.097, 0.2] mg), high dispersed (0.162 [0.113, 0.233] mg), and low (0.115 [0.077, 0.172] mg)—produced significantly more sugar than the control group (0.036 [0.019, 0.069] mg), with no significant differences among the damage treatments (Fig. 1b). Finally, the proportion of secreting nectaries was also significantly affected by leaf damage (Table 1), with only the high dispersed damage treatment (0.527 [0.454, 0.599]) differing significantly from the control (0.288 [0.226, 0.359]), as well as from the low (0.378 [0.312, 0.449]) and high concentrated (0.355 [0.294, 0.421]) damage treatments (Fig. 1c). Statistics for Tukey pairwise comparisons for all nectar responses are presented in Table S1 A-C, of the supplementary material.

**Figure 1.**
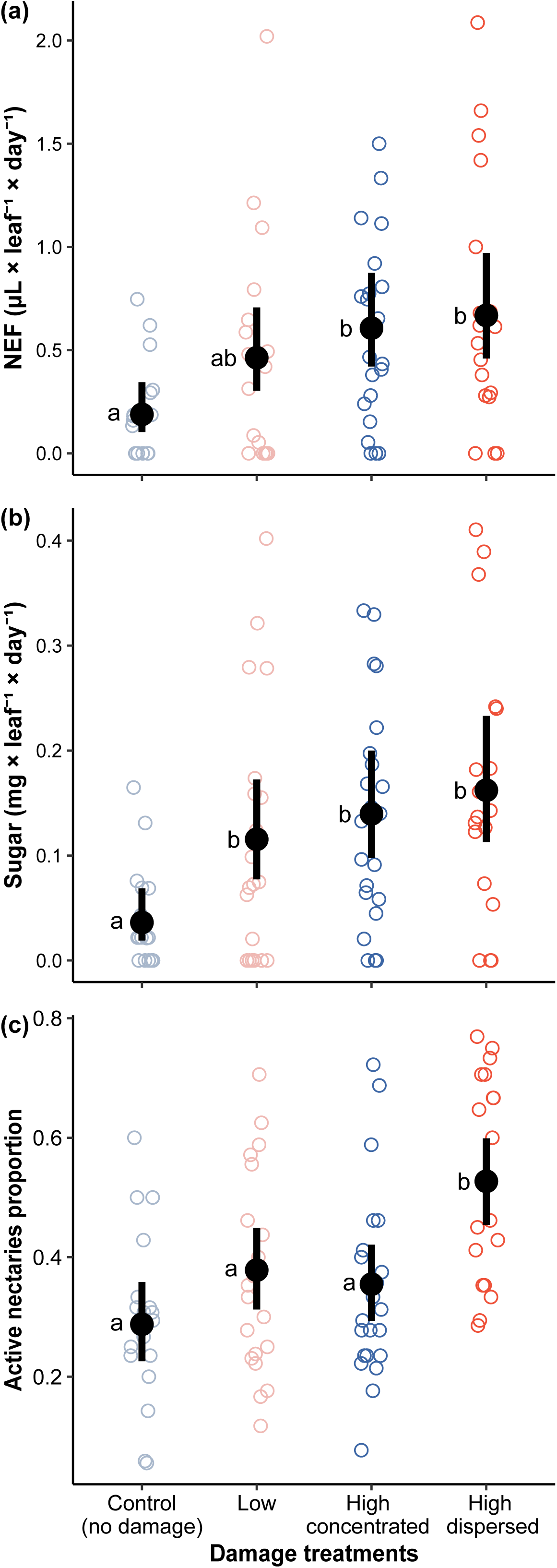
EFN volume production and sugar content for leaf 2 (damaged leaf, except for control plants) of wild cotton (*Gossypium hirsutum*) plants under different leaf damage treatments. Panels show: (a) per-nectary EFN volume, (b) per-nectary EFN sugar content, and (c) the proportion of active nectaries per plant. Black dots and error bars indicate EMMs and 95% CIs based on GLMMs. Single values are plotted (open colour circles). Different letters indicate significant differences between treatments according to Tukey’s post hoc test (p < 0.05).

**Table 1.** Wald χ² tests for the GLMMs assessing the effects of leaf damage treatments and plant height on EFN (volume, total sugar, and proportion of active nectaries) and ant activity (patrolling, and attack on caterpillars).

| Response variable | Effect | df | $\chi^2$ | *P* |
| --- | --- | --- | --- | --- |
| EFN volume | Damage treatment | 3 | 13.69 | **0.0003** |
|  | Plant height | 1 | 1.41 | 0.235 |
| Sugar production | Damage treatment | 3 | 16.95 | **0.0007** |
|  | Plant height | 1 | 0.21 | 0.6394 |
| Proportion of nectaries active | Damage treatment | 3 | 23.39 | **< 0.0001** |
|  | Plant height | 1 | 1.246 | 0.2643 |
| Patrolling ants | Damage treatment | 3 | 14.99 | **0.0018** |
|  | Plant height | 1 | 3.12 | 0.0773 |
| Attack on caterpillars | Damage treatment | 3 | 10.92 | **0.0121** |
|  | Plant height | 1 | 1.53 | 0.2454 |

### Ant recruitment and anti-herbivore defensive response

We identified 10 ant species on the experimental cotton plants, with *Monomorium ebeninum*, *Brachymyrmex australis*, and *Forelius pruinosus* being the most abundant (Table S2, Supplementary Material). The leaf damage treatment significantly affected ant abundance (Table 1). Specifically, plants subjected to the high dispersed damage treatment attracted significantly more ants (5.45 [3.86, 7.7] ants) than controls (2.33 [1.61, 3.35] ants) and plants exposed to high concentrated damage (2.45 [1.69, 3.55] ants). The low damage treatment (3.01 [2.07, 4.36] ants) did not differ significantly from either the control or the high damage treatments (Fig. 2a; Table S1). Consistently, caterpillars on plants with high dispersed damage in turn had the highest probability of being attacked by ants (0.886 [0.713, 0.961]), which was significantly greater than the probability of attack on controls (0.442 [0.27, 0.629]) and plants with low damage (0.499 [0.293, 0.705]) treatments. The attack probability for the high concentrated damage treatment (0.629 [0.429, 0.794]) exhibited an intermediate value and did not differ significantly from controls or the other damage treatments (Fig. 2b; Table S1).

**Figure 2.**
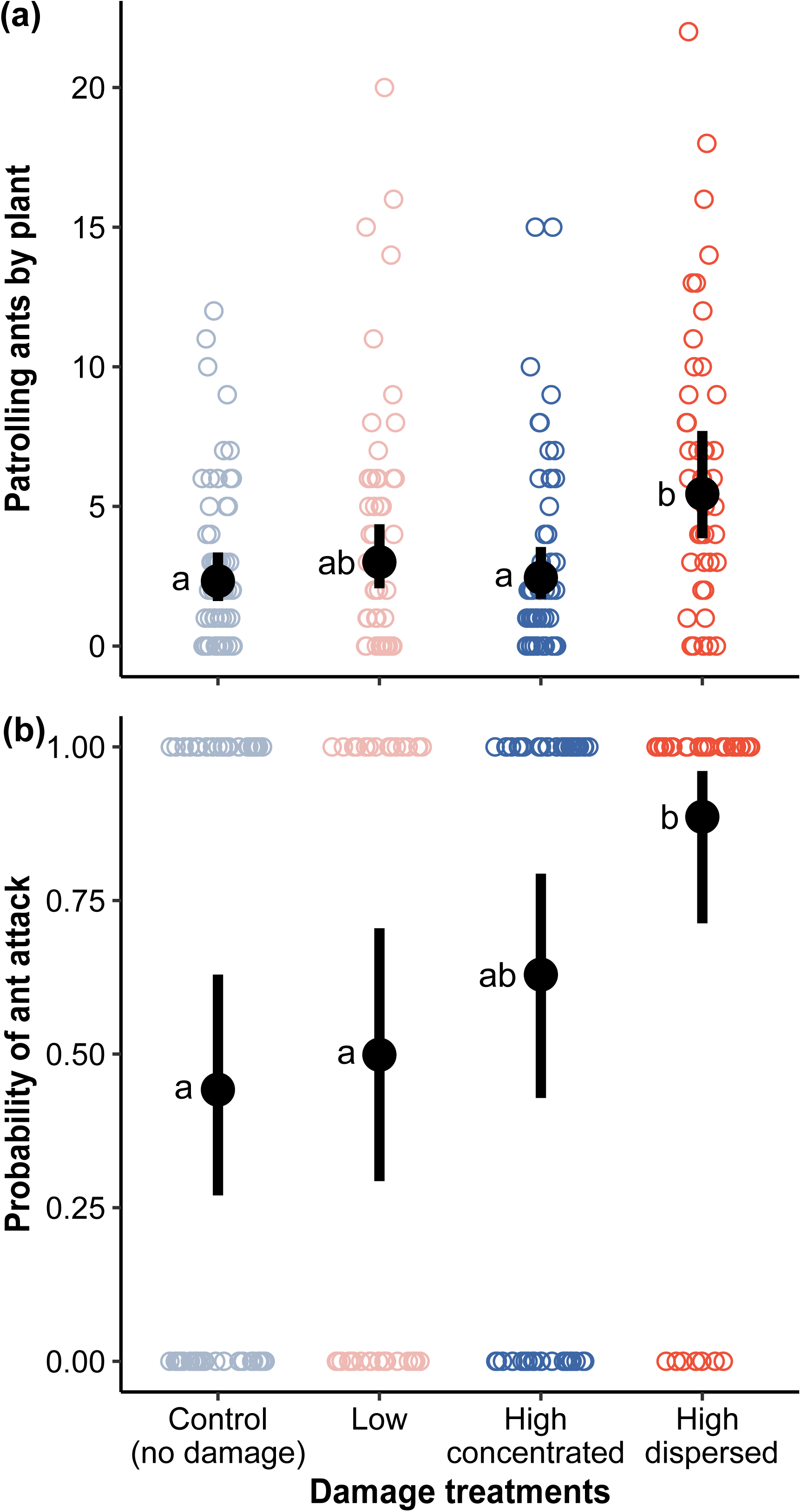
Ant activity on wild cotton (*Gossypium hirsutum*) plants under different leaf damage treatments. Panels: (a) mean number of patrolling ants per plant, and (b) probability of ant attack on caterpillars (i.e., indirect defence bioassays). Black dots and error bars indicate EMMs and 95% CIs based on GLMMs. Single values are plotted (open colour circles). Different letters indicate significant differences between treatments according to Tukey’s post hoc test (p < 0.05).

## Discussion

Effects of damage amount on wild cotton EFN induction varied as a function of the trait measured, with low damage significantly inducing (relative to controls) EFN sugar content and behaving similarly to the high damage treatments, but did not induce either EFN volume or the proportion of active nectaries. Within-plant dispersion of damage, on the other hand, had no detectable effects on nectar volume or sugar content (i.e., no difference between concentrated vs. dispersed treatments), but did have a clear effect on the proportion of secreting nectaries whereby only plants in the high dispersed damage group differed significantly from controls as well as from the other damage groups, whereas the latter did not differ from controls. Consistently, dispersed damage was the only treatment to significantly increase ant abundance and the probability of attack relative to controls. Overall, these findings suggest that variation in the spatial uniformity of leaf damage within the plant (i.e., concentrated vs. dispersed damage), rather than damage amount, is the main driver of ant recruitment and indirect defence via EFN induction. These results highlight a previously underappreciated role of variability in the spatial uniformity of herbivore damage in shaping ant-plant mutualisms and call for greater attention to how herbivory patterns influence the induction of plant defences.

### Extrafloral nectar

Leaf damage amount has been shown to correlate positively with EFN induction in other plant species (Rios et al., 2008; Xu et al., 2014; Calixto et al., 2021), suggesting that plants are able to allocate to induced responses proportionally to perceived herbivore threats (e.g., as a function of herbivory severity). Our findings provide partial support for this. Both high damage treatments, but not the low damage treatment, resulted in significantly more EFN volume secreted compared to controls, suggesting that damage amount matters for this trait. In contrast, low damage increased (significantly) total sugar production to a similar extent relative to the high damage treatments, i.e., the induction of this trait was not contingent on damage amount. This latter result is similar to that observed, for example, in the shrub *Banisteriopsis malifolia*, for which leaf damage increases EFN sugar secretion to similar degree regardless of damage severity (Pimenta et al., 2025). In particular, previous work with wild cotton found that EFN volume induction behaved similarly across levels of leaf damage (Abdala-Roberts et al., 2019b), albeit involving overall lower damage levels (relative to the present study) and the combination of mechanical damage with oral secretions from the generalist *Spodoptera frugiperda*. However, in a recent study using natural herbivory by *A. argillacea* we found that low herbivory significantly induced EFN volume (Ramírez-Delgado et al., 2025). This suggests that differences in the induction of EFN volume between natural (but contingent on the caterpillar species) and mechanical damage can have an important influence on EFN induction. In that same study, however, low herbivory did not induce EFN sugar content, which together with our present results, suggests that the induction of sugar content behaves similarly irrespective of damage amount, i.e., EFN traits are not induced equally by different degrees of herbivory severity and based on whether natural vs. mechanical damage is applied. Further experimentation is needed to assess better the role of herbivory amount on wild cotton, such as the inclusion of a broader range and number of damage levels combined with mechanical vs. natural damage treatments, to better understand herbivory amount’s effect on the induction of multiple EFN-associated traits.

Our results also revealed that within-plant variability in the spatial uniformity of leaf damage significantly affected EFN induction, specifically by altering the proportion of secreting nectaries. Contrary to predictions, plants subjected to high dispersed damage were the only group that differed significantly from controls and other damage treatments for this EFN response. The fact that recent work shows that cotton plants are less able to compensate for concentrated than dispersed damage (Ramírez-Delgado et al., 2025), suggests that EFN induced responses are not aligned with differential resource reallocation costs from contrasting herbivory patterns but rather respond to other factors such as ant foraging patterns. We come back to this later on and next speculate on two proximal, non-exclusive induction-related processes. First, high dispersed damage presumably locally induced a greater number of nectaries (i.e., by spreading damage across more leaves per plant, and despite lower damage amount per leaf), while maintaining similar EFN production per nectary compared to the other damage treatments (see also Wäckers et al., 2001). Second, dispersed damage may have also triggered a stronger systemic response, leading to increased nectar production in undamaged leaves. Supporting the latter, we observed that the dispersed damage treatment was the only one for which all plants exhibited nectar secretion in at least two non-damaged leaves, whereas for low damage and concentrated damage nectar induction was restricted to damaged leaves for most plants. Teasing apart the relative contributions of EFN local vs. systemic induced responses would be a worthwhile goal in future work, including also a more detailed investigation of physiological mechanisms driving systemic EFN induced responses to dispersed damage in a defensive trait previously shown to be preeminently shaped by local induction patterns (Abdala-Roberts et al., 2019b). These may include, for example, within-plant volatile signalling, i.e., among stems or leaves (Hernández-Zepeda et al., 2018) or signal transport through the xylem, which may reach undamaged leaves and upregulate nectar production (Chatt et al., 2021). However, these explanations are merely speculative at this stage. Finally, considering the role of herbivore chemical elicitors using natural damage by different insect herbivores would also be useful, as even though EFN responses are often strongly determined by mechanical damage per se, herbivore-specific factors merit attention in studying the mechanisms of local vs. systemic induction of EFN under contrasting herbivory patterns.

### Ant abundance and anti-herbivore response

The ant species recorded during the field experiment have been previously reported on wild cotton, though involving different species composition and relative abundances in some cases. For example, Reyes-Hernández et al., (2022) reported high abundances of *Crematogaster* on wild cotton in a tropical dry forest gap, while Vázquez-Barrios et al., (2021) found that *Camponotus planatus* and *Paratrechina longicornis* were dominant species in a coastal scrubland. In contrast, our study recorded high abundances of *Monomorium ebeninum*, *Brachymyrmex australis*, and *Forelius pruinosus*. Although these small-bodied species are not strictly predatory (Díaz-Castelazo et al., 2004), their persistent presence on plants may discourage herbivores through disturbance or harassment, potentially leading to reduced overall damage (Bentley, 1977; Rudgers et al., 2003). Larger, more aggressive species, including predatory ants in *Pseudomyrmex* and *Camponotus*, were much less abundant but have been previously documented on wild cotton at coastal sites (Vázquez-Barrios et al., 2021). Ant competitive dynamics and the availability of alternative plant resources or prey may influence ant community composition (Byk & Del-Claro, 2011; Nogueira et al., 2015) and warrant further examination across wild cotton populations.

Mirroring effects on the proportion of secreting nectaries, ant abundance (pooled across species) was highest on plants subjected to high dispersed damage. This was likely due to an increased availability of secreting nectaries per plant under this treatment, enhancing ant recruitment and patrolling activity. Consistent with this finding, attack rates on caterpillars were highest on plants subjected to dispersed damage, suggesting increased ant abundance led to higher attack rates, i.e., a density-mediated effect (Abdala-Roberts et al., 2012). It is also possible that dispersed damage increased ant aggressiveness or foraging efficiency, independently of ant numbers, as previous work in other systems has reported that plants influence ant behaviour through differences in nectar composition (Ness, 2003; Ness et al., 2006; Shenoy et al., 2012), i.e., a trait-mediated effect (Abdala-Roberts & Mooney, 2015). Regardless of the mechanism, and while this finding suggests dispersed damage could lead to heightened indirect defence, it is important to note that recorded attacks on caterpillars may not be representative of actual ant effectiveness in reducing herbivory. In addition, herbivory levels are highly fluctuating and are overall often low at cotton populations (Abdala-Roberts et al., 2019a), suggesting that the benefits to plant fitness of short-term ant responses may only accrue under a specific set of conditions (e.g., periods of high and consistent damage over several weeks and ant communities dominated by more aggressive species). It is important to also note that although using dead caterpillars has proven to be a useful as well as realistic method for testing ant predatory or deterrence behaviour against insect herbivores (Grof-Tisza et al., 2024), this approach may underestimate attack rates (as live caterpillars may move and attract more attention) or overestimate successful deterrence and linkages to indirect defence on plants as live caterpillars can actively defend themselves (Lövei & Ferrante, 2017; Nagy et al., 2020). Within this context, an important challenge will be to disentangle ant-plant-herbivore interaction mechanisms via experimental procedures such as ant exclusions across sites with contrasting herbivore and ant abundances and composition, and testing for such effects over longer periods of time during the growing season.

### An ecological-based mechanism linking cotton direct and indirect defences in response to herbivory variability

The implications of our results can be interpreted within the broader context of cotton’s extensive defensive arsenal, including direct chemical defences. Concentrated herbivory typically imposes greater fitness costs on plants (Marquis, 1992; Wetzel et al., 2023), a notion that is consistent with recent findings with wild cotton (Ramírez-Delgado et al., 2025). We predicted that direct chemical defences (e.g., phenolics, terpene aldehydes) should be more strongly induced under such attacks. If this is the case, in theory, it would not only reduce herbivory but also promote its spread from leaf-to-leaf, further contributing to dampen the costs of localised damage (Marquis, 1996; Karban & Yang, 2020). Once damage becomes dispersed, the plant’s strategy can shift to enhancing indirect defence, aligning with our results of increased EFN production and ant recruitment. Based on this, a selective feedback loop could be in place whereby selection for direct and indirect defences becomes entrained: the induction of chemical defences against concentrated attacks is reinforced because it leads to dispersed damage, which in turn promotes stronger or more effective EFN-mediated ant defence. Current research in our group is aimed at testing the effects of variability in the spatial uniformity of herbivore damage on both chemical defences and EFN, as well as assessing the fitness costs of different herbivory spatial uniformity patterns. From this, we can then expand and evaluate these responses across populations that experience contrasting ecological conditions (e.g., varying ant composition, herbivore pressure, and abiotic stress). In closing, our results underscore the complexity of plant-insect multitrophic interactions in natural ecosystems and emphasise the need to understand how plants optimise their defensive investments across varying herbivory contexts in order to maximise the benefits of indirect defence derived from mutualistic interactions with predators.

