## Supplemental information for "From leaves to defenders: how the amount and dispersion of leaf damage affect extrafloral nectar production and ant-mediated protection in wild cotton"

### Table of Contents:


| | |
|---|---|
| **Figure S1** | Page 2 |
| **Table S1** | Page 4 |
| **Table S2** | Page 5 |

(a)

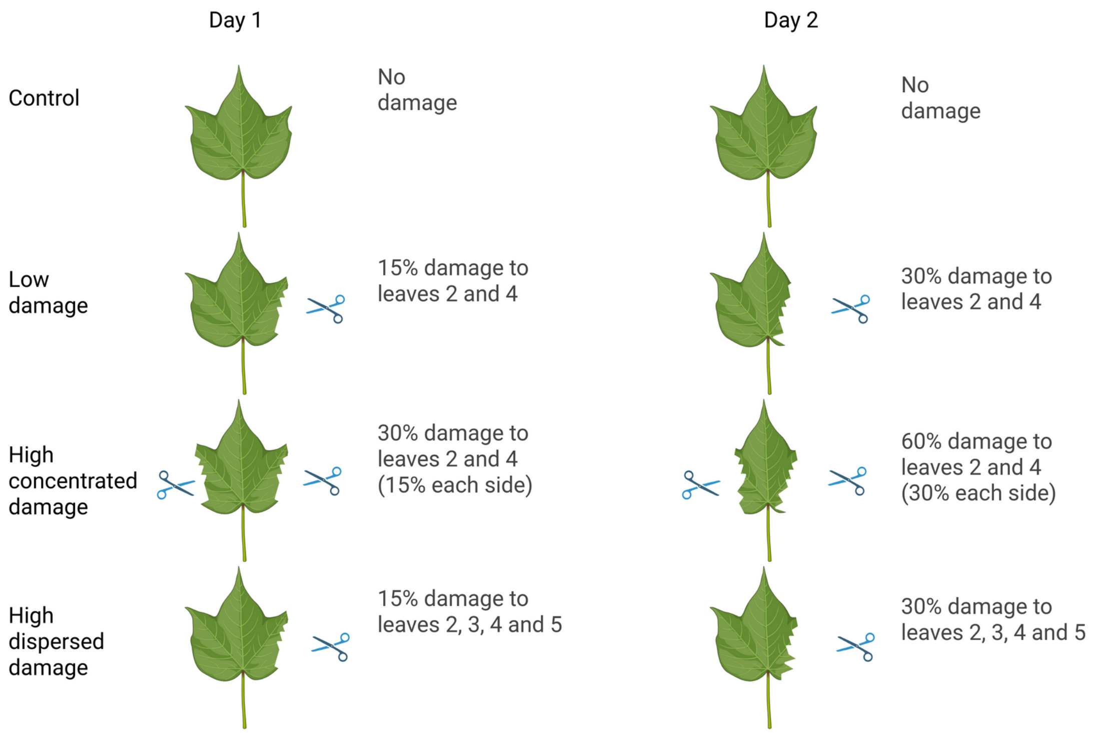


(b)

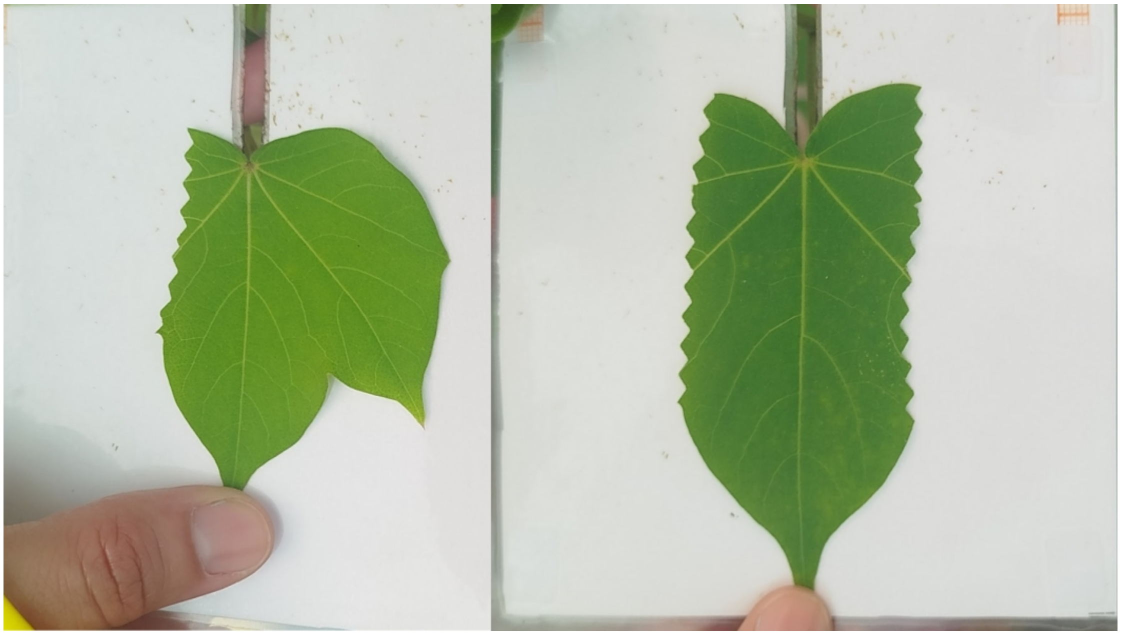

**Figure S1.** (a) Amount of leaf tissue removed each day during the damage application period (2 days). (b) Examples of leaf damage treatments after two days of damage. Left: a leaf with one side removed; Right: a leaf with both sides removed. Panel (a) created in BioRender. https://BioRender.com/omjj04e

**Table S1**. Contrasts of treatments using Tukey tests. Treatments: control, low damage (LD), high damage concentrated (HDC), and high damage dispersed (HDD). Bold P-values < 0.05.

| Contrast | Ratio | SE | df | Z ratio | P-value |
|---|---|---|---|---|---|
| A) Nectar volume | | | | | |
| Control / LD | 0.43 | 0.162 | ∞ | -2.246 | 0.1111 |
| Control / HDC | 0.334 | 0.12 | ∞ | -3.045 | **0.0125** |
| Control / HDD | 0.283 | 0.102 | ∞ | -3.487 | **0.0027** |
| LD / HDC | 0.776 | 0.221 | ∞ | -0.891 | 0.8097 |
| LD / HDD | 0.659 | 0.189 | ∞ | -1.458 | 0.4634 |
| HDC / HDD | 0.849 | 0.226 | ∞ | -0.616 | 0.927 |
| B) Total sugar | | | | | |
| Control / LD | 0.313 | 0.1213 | ∞ | -2.997 | **0.0145** |
| Control / HDC | 0.259 | 0.0978 | ∞ | -3.578 | **0.002** |
| Control / HDD | 0.223 | 0.0832 | ∞ | -4.02 | **0.0003** |
| LD / HDC | 0.827 | 0.2259 | ∞ | -0.696 | 0.8985 |
| LD / HDD | 0.712 | 0.1975 | ∞ | -1.225 | 0.611 |
| HDC / HDD | 0.861 | 0.2268 | ∞ | -0.568 | 0.9417 |
| C) Proportion of active nectaries | | | | | |
| Control / LD | 0.664 | 0.1506 | ∞ | -1.807 | 0.2697 |
| Control / HDC | 0.734 | 0.1618 | ∞ | -1.401 | 0.4987 |
| Control / HDD | 0.362 | 0.0795 | ∞ | -4.625 | **<.0001** |
| LD / HDC | 1.107 | 0.2249 | ∞ | 0.5 | 0.9591 |
| LD / HDD | 0.546 | 0.1172 | ∞ | -2.818 | **0.025** |
| HDC / HDD | 0.493 | 0.1029 | ∞ | -3.388 | **0.0039** |
| D) Number of ants | | | | | |
| Control / LD | 0.774 | 0.195 | ∞ | -1.017 | 0.7397 |
| Control / HDC | 0.95 | 0.244 | ∞ | -0.201 | 0.9971 |
| Control / HDD | 0.427 | 0.106 | ∞ | -3.425 | **0.0034** |
| LD / HDC | 1.227 | 0.321 | ∞ | 0.783 | 0.862 |
| LD / HDD | 0.551 | 0.139 | ∞ | -2.357 | 0.0855 |
| HDC / HDD | 0.449 | 0.113 | ∞ | -3.194 | **0.0077** |
| E) Ant attack | | | | | |
| Control / LD | 0.796 | 0.4671 | ∞ | -0.388 | 0.9801 |
| Control / HDC | 0.467 | 0.2706 | ∞ | -1.314 | 0.5538 |
| Control / HDD | 0.102 | 0.0727 | ∞ | -3.197 | **0.0076** |
| LD / HDC | 0.586 | 0.3677 | ∞ | -0.852 | 0.8297 |
| LD / HDD | 0.128 | 0.0947 | ∞ | -2.776 | **0.0282** |
| HDC / HDD | 0.218 | 0.1486 | ∞ | -2.235 | 0.1138 |

37 **Table S2**. Number of plants on which a given ant species was recorded and the total number of

38 ants recorded per species per day.

| | Day 1 | | Day 2 | |
|---|---|---|---|---|
| Species | Plants patrolled | Total ants | Plants patrolled | Total ants |
| *Monomorium ebeninum* | 30 | 139 | 32 | 156 |
| *Brachymyrmex australis* | 26 | 120 | 32 | 122 |
| *Forelius pruinosus* | 25 | 96 | 26 | 130 |
| *Pheidole* sp. | 6 | 9 | 4 | 8 |
| *Solenopsis geminata* | 4 | 4 | 2 | 3 |
| *Crematogaster torosa* | 2 | 4 | 2 | 4 |
| *Nylanderia steinheili* | 2 | 2 | 1 | 1 |
| *Pseudomyrmex elongatulus* | 2 | 2 | 1 | 1 |
| *Camponotus planatus* | 1 | 2 | - | - |

39
